# Trace levels of peptidoglycan in serum underlie the NOD-dependent cytokine response to endoplasmic reticulum stress

**DOI:** 10.1101/547885

**Authors:** Raphael Molinaro, Robert Flick, Dana J. Philpott, Stephen E. Girardin

**Author notes:** To whom correspondence should be addressed: Stephen E. Girardin, Ph.D.

## Abstract

NOD1 and NOD2 are intracellular sensors of bacterial peptidoglycan that belong to the Nod-like receptor (NLR) family of innate immune proteins. In addition to their role as direct bacterial sensors, it was recently proposed that NOD proteins could detect endoplasmic reticulum (ER) stress induced by thapsigargin, an inhibitor of the sarcoplasmic or endoplasmic reticulum calcium ATPase family (SERCA) that pumps Ca^2+^ into the ER, resulting in pro-inflammatory signalling. Here, we confirm that thapsigargin induces pro-inflammatory signalling in epithelial cells in a NOD-dependent manner. However, the effect was specific to thapsigargin, as tunicamycin and the subtilase cytotoxin SubAB from Shiga toxigenic *Escherichia coli,* which induce ER stress by other mechanisms, did not induce cytokine expression. The calcium ionophore A23187 also induced NOD-dependent signalling, and the calcium chelator BAPTA-AM blunted thapsigargin-dependent pro-inflammatory signalling, showing NOD proteins responded to a rise in intracellular Ca^2+^. Since intracellular Ca^2+^ directly affects vesicular trafficking, we tested if thapsigargin-induced NOD activation required endocytosis. Our results demonstrate that both endocytosis and the addition of serum to the cell culture medium were required for thapsigargin-mediated NOD activation. Finally, we analyzed cell culture grade fetal calf serum as well as serum from laboratory mice by high-pressure liquid chromatography and mass spectrometry, and identified the presence of various peptidoglycan fragments. We propose that cellular perturbations that affect intracellular Ca^2+^ can trigger internalization of peptidoglycan trace contaminants found in culture serum, thereby stimulating pro-inflammatory signalling. The presence of peptidoglycan in animal serum suggests that a homeostatic function of NOD signalling may have been previously overlooked.

## Introduction

Detection of microbes by the innate immune system relies on several families of pattern recognition molecules (PRMs) that recognize conserved microbe-associated molecular patterns (MAMPs) that are highly conserved and are not produced by the non-infected host. Among those families of PRMs are the Toll-like receptors (TLRs) and Nod-like receptors (NLRs). In addition to the detection of MAMPs, certain NLR proteins, such as NLRP3, detect cellular perturbations or molecules, known as danger-associated molecular patterns (DAMPs) that can arise as a result of an infection or following aseptic tissue damage (1).

NOD1 and NOD2 are two members of the NLR family of PRM that detect bacterial peptidoglycan (2). The specificity of NOD1 and NOD2 for peptidoglycan motifs is extremely high, as NOD1 detects *meso*-diaminopimelic acid-containing N-acetyl muramic acid (MurNAc)-tripeptide (Mur-TriDAP) found predominantly in Gram-negative bacteria (3–6), whereas NOD2 detects muramyl dipeptide (MDP) found in both Gram-negative and Gram-positive bacteria (3,7,8). Detailed studies on the minimal structural requirements of the peptidoglycan ligands needed for NOD1 or NOD2 activation revealed that the MurNAc moiety is not required for NOD1 activation as the D-Glu-*meso*-DAP dipeptide (iE-DAP) is sufficient for detection and innate immune activation by this PRM (3,9). NOD2, on the other hand, can only be activated by muramyl dipeptides that have an intact MurNAc ring structure, and the sugar has to be attached to a dipeptide moiety (L-Ala-D-Glu or L-Ala-D-isoGln) (3,10). Importantly, studies have shown that both NOD1 and NOD2 can directly bind to TriDAP and MDP, respectively, thus showing that NOD1 and NOD2 are *bona fide* cytoplasmic receptors (11–13), and that this interaction requires the leucine - rich repeat region of NOD1 and NOD2 proteins (14,15).

While functional and binding studies demonstrate that NOD proteins have an extreme specificity for certain peptidoglycan fragments that is conserved in multiple Vertebrates species, recent studies have suggested that NOD signaling could also be triggered by viral infection (16), small Rho GTPases regulating the cytoskeleton (17), and endoplasmic reticulum (ER) stress (18), implying that peptidoglycan-independent mechanisms of NOD stimulation may exist. This suggests that in addition to their high specificity towards peptidoglycan fragments, NOD1 and NOD2 may serve as promiscuous sensors of multiple and unrelated cellular stresses. Here, we aimed to better characterize how NOD proteins trigger pro-inflammatory signaling in response to ER stress. While we confirm that thapsigargin, a specific inhibitor of the ER sarcoplasmic or endoplasmic reticulum calcium ATPase family (SERCA) calcium pump, triggers pro-inflammatory signalling in a NOD-dependent manner (18), our results suggest that this effect is actually mediated by the Ca^2+^-dependent internalization of peptidoglycan trace fragments found in the fetal calf serum added to cell culture media. Of note, low levels of peptidoglycan in human serum have already been reported, and were shown to stimulate hyphal growth of *Candida albicans* (19). Together, our observations suggest that cellular perturbations that lead to increased intracellular Ca^2+^ levels may inadvertently trigger NOD-dependent signalling through the internalization of peptidoglycan contaminants, which offers an alternative explanation for the proposed promiscuous activation of NOD receptors by multiple unrelated stresses. These observations also suggest that chronic homeostatic peptidoglycan sensing by NOD proteins may impact multiple cellular processes in ways that have been overlooked previously and open up interesting questions in innate immunity, relating to understanding the physiological role of circulating peptidoglycan at homeostasis, both at the cellular and the tissue level.

## Results and Discussion

The human intestinal epithelial cell line HCT116, either wild type (WT) or knockout out through CRISPR-Cas9 for NOD1, NOD2 or both NOD1 and NOD2 (NOD 1/2 double knockout or DKO) described previously (20) were stimulated with the ER stress inducer thapsigargin, which inhibits the SERCA pump that transports Ca^2+^ to the ER lumen. While thapsigargin induced ER stress similarly in the four cell lines tested, as determined by the transcriptional upregulation of the heat shock protein GRP78 (also known as BiP and encoded by the *HSPA5* gene) (Fig. 1A), transcriptional upregulation of the pro-inflammatory cytokines CXCL1 (Fig. 1A), IL-8 (Fig. S1A) as well as the chemokine CCL20 (Fig. S1A) was significantly blunted in NOD2 KO and NOD1/2 DKO HCT116 cells, in line with previous results (18). In NOD1 KO cells, a non-significant trend for reduced expression of CXCL1 and IL-8 was also noticed following thapsigargin stimulation, and significant reduction of CCL20 of expression was observed (Figs. 1A and S1A). Together, these results suggest that, in HCT116 cells, NOD2 and to a lesser extent NOD1, contribute to the upregulation of pro-inflammatory signalling induced by thapsigargin. On average, NOD1/2 DKO cells displayed an 8.35 fold reduction in CXCL1 induction following thapsigargin stimulation, a 6.33 fold reduction in IL-8 expression, and a 2.85 fold reduction in CCL20 expression.

**Figure 1.**
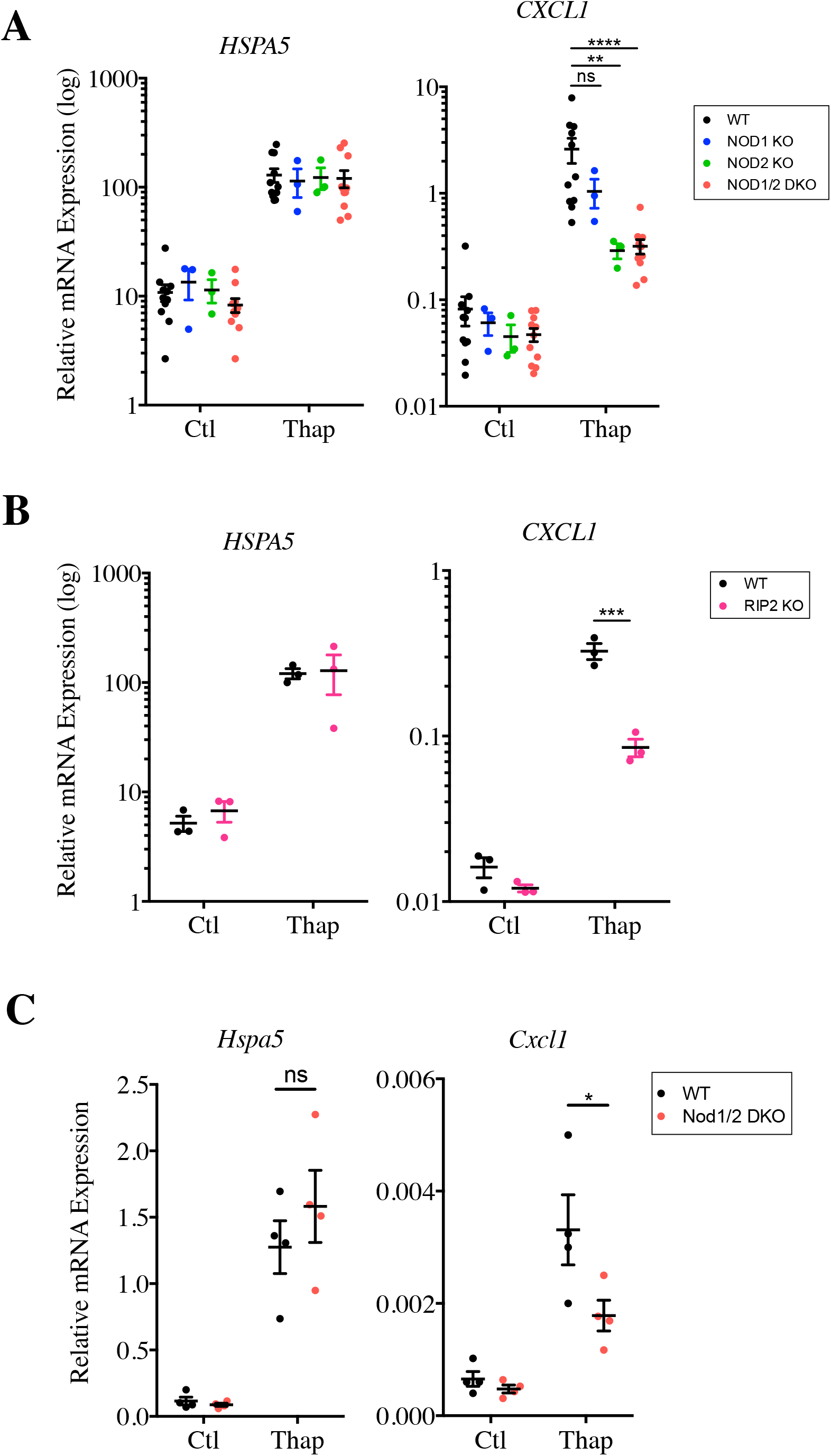
Thapsigargin induces pro-inflammatory cytokine expression in a NOD-dependent manner. **(A-B)** Expression of *HSPA5* and *CXCL1* in wild type (WT), NOD1 knockout (KO), NOD2 KO and NOD1/2 double KO (DKO) HCT116 cells **(A)** or WT and RIP2 KO HCT116 cells **(B)** following stimulation with 0.1*μ*g/ml thapsigargin for 4 hours measured by qPCR. **(C)** Expression of *Hspa5* and *Cxcl1* in primary intestinal organoids from wild type (WT), and NOD1/2 double KO (DKO) mice following stimulation with 1*μ*g/ml thapsigargin for 4 hours measured by qPCR. Each point is from an independent experiment and is the average of three technical replicates. *, ** and ****, p < 0.05, 0.01 and 0.0001, respectively. NS, not significant.

To demonstrate that this effect requires the NOD adaptor protein RIP2, we engineered HCT116 RIP2 KO cells by targeting the *RIPK2* gene (encoding RIP2) using CRISPR-Cas9. As expected, RIPK2-targeted cells were insensitive to the synthetic NOD ligands MDP and iE-DAP in an NF-κB luciferase assay (Fig. S2A). Similar to NOD1/2 DKO cells, RIP2 KO HCT116 cells displayed normal induction of ER stress-dependent BiP/GRP78/HSPA5 (Fig. 1B), but significantly reduced upregulation of CXCL1 (Fig. 1B), IL-8 and CCL20 (Fig. S1B). On average, RIP2 KO cells displayed a 3.82 fold reduction in CXCL1 induction following thapsigargin stimulation, a 13.48 fold reduction in IL-8 expression, and a 3.04 fold reduction in CCL20 expression. To further validate our findings, we reproduced these results with other clones of NOD1/2 DKO and RIP2 KO cells (Fig. S2B-C). Finally, we isolated primary intestinal organoids from WT and Nod1/2 DKO mice and stimulated those with thapsigargin. In previous work, we identified using RNAseq that thapsigargin potently stimulated cytokine expression in murine organoids (21). Similar to our results with the human intestinal cell line HCT116, we observed that while thapsigargin triggered comparable expression of BiP/Hspa5 in WT and Nod1/2 DKO organoids, Cxcl1 expression was significantly blunted in Nod1/2 DKO organoids (Fig. 1C), although the effect was not as drastic as in HCT116 cells, suggesting that NOD-dependent signalling only represents a subset of thapsigargin-dependent pro-inflammatory cascades in murine organoids. Together, these results confirm the previous findings (18) that NOD1 and NOD2 are critical for inflammatory signalling induced by thapsigargin.

We next aimed to define if other ER stress inducers trigger pro-inflammatory signalling in a NOD-dependent manner in HCT116 cells. We first used tunicamycin, a molecule that induces ER stress by preventing protein glycosylation in the ER lumen, thereby provoking the accumulation of misfolded proteins in the ER. This mechanism of ER stress induction is distinct from the one triggered by thapsigargin, which relies on the inhibition of Ca^2+^ accumulation in the ER lumen. Interestingly, while tunicamycin induced potent upregulation of BiP/HSPA5 expression in WT, NOD1 KO, NOD2 KO and NOD1/2 DKO HCT116 cells (Fig. 2A), it did not stimulate expression of CXCL1 or IL8 (Fig. 2A), suggesting that induction of pro-inflammatory signalling by thapsigargin is likely caused by the effect of the inhibitor on intracellular Ca^2+^ levels rather than caused by ER stress *per se*. To confirm these findings, we induced ER stress by a third mechanism, namely the targeting and degradation of the BiP protein by the SubAB toxin of Shiga toxigenic *Escherichia coli* (STEC), which causes ER stress as a consequence of reduced levels of luminal BiP chaperone (22,23). SubAB strongly induced upregulation of BiP/HSPA5 expression in WT, NOD1 KO, NOD2 KO and NOD1/2 DKO HCT116 cells, while a mutant toxin unable to cleave BiP was unable to do so (Fig. 2B). However, and similar to tunicamycin, SubAB did not trigger expression of CXCL1 or IL8 (Fig. 2B), thus further demonstrating that, at least in HCT116 cells, ER stress does not induce NF-κB-dependent pro-inflammatory cytokines such as CXCL1 and IL8, and that thapsigargin action on pro-inflammatory signalling is likely caused by its capacity to rise intracellular Ca^2+^ levels.

**Figure 2.**
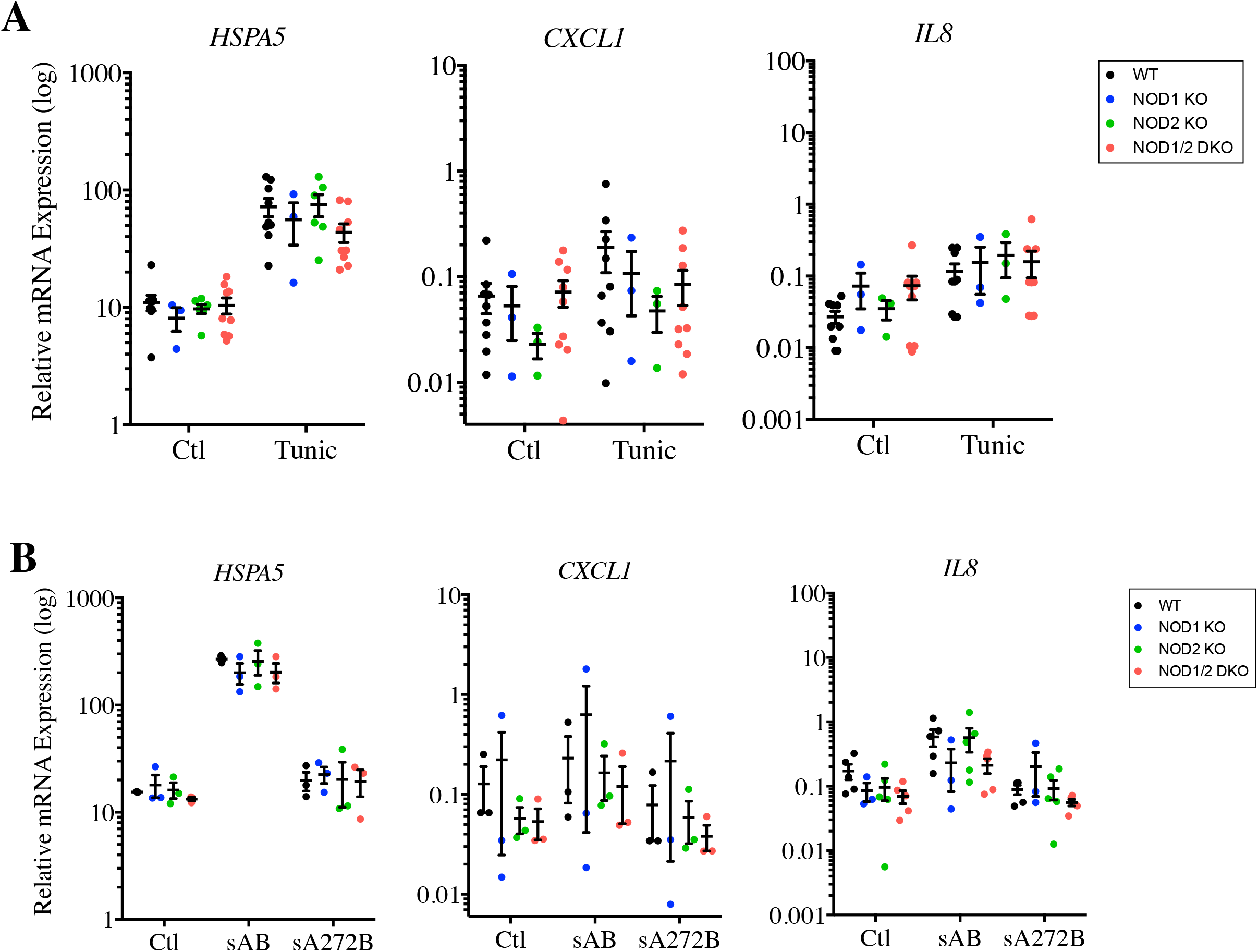
ER stress inducers that do not directly target Ca^2+^ stores do not trigger NOD-dependent pro-inflammatory cytokine expression. **(A-B)** Expression of *HSPA5, CXCL1* and *IL8* in wild type (WT), NOD1 knockout (KO), NOD2 KO and NOD1/2 double KO (DKO) HCT116 cells following stimulation with 1*μ*g/ml tunicamycin **(A)** or 50 ng/ml SubAB toxin from Shiga toxigenic *Escherichia coli,* either wild type (sAB) or mutated on position 272 (sA272B), which abolishes its activity, for 4 hours **(B)** and measured by qPCR. Each point is from an independent experiment and is the average of three technical replicates.

To directly test the role of intracellular Ca^2+^ levels in NOD-dependent pro-inflammatory signalling, WT and NOD1/2 DKO cells were stimulated with A23187, a Ca^2+^ ionophore that causes massive influx of Ca^2+^ into the cytosol. As expected, stimulation with A23187 caused upregulation of BiP/HSPA5 in both WT and NOD1/2 DKO cells (Fig. 3A). Similarly to thapsigargin, A23187 also stimulated expression of CXCL1 and IL8, which was significantly blunted in NOD1/2 DKO cells (Fig. 3A), thus showing that the rise intracellular Ca^2+^ levels caused NOD-dependent pro-inflammatory signalling. Moreover, using BAPTA-AM, a cell-permeant Ca^2+^ chelator, we directly tested the role of intracellular Ca^2+^ levels in thapsigargin-stimulated cells. While BAPTA-AM did not significantly affect ER stress induced by thapsigargin (Fig. 3B), in line with the fact that BAPTA-AM does not prevent thapsigargin-dependent depletion of luminal ER stores of Ca^2+^ but neutralizes the effects of cytosolic Ca^2+^, it blunted the induction of CXCL1 and IL8 in WT cells but not in NOD1/2 DKO cells. As a result, thapsigargin-induced expression of pro-inflammatory cytokines was not significantly dependent on NOD1/2 in BAPTA-AM-treated cells, although a trend for decreased expression was still observed in NOD1/2 DKO cells, possibly because BAPTA-AM was unable to fully neutralize the effects of intracellular Ca^2+^. Together, we conclude that the rise of intracellular Ca^2+^ levels caused NOD-dependent pro-inflammatory signalling in HCT116 cells.

**Figure 3.**
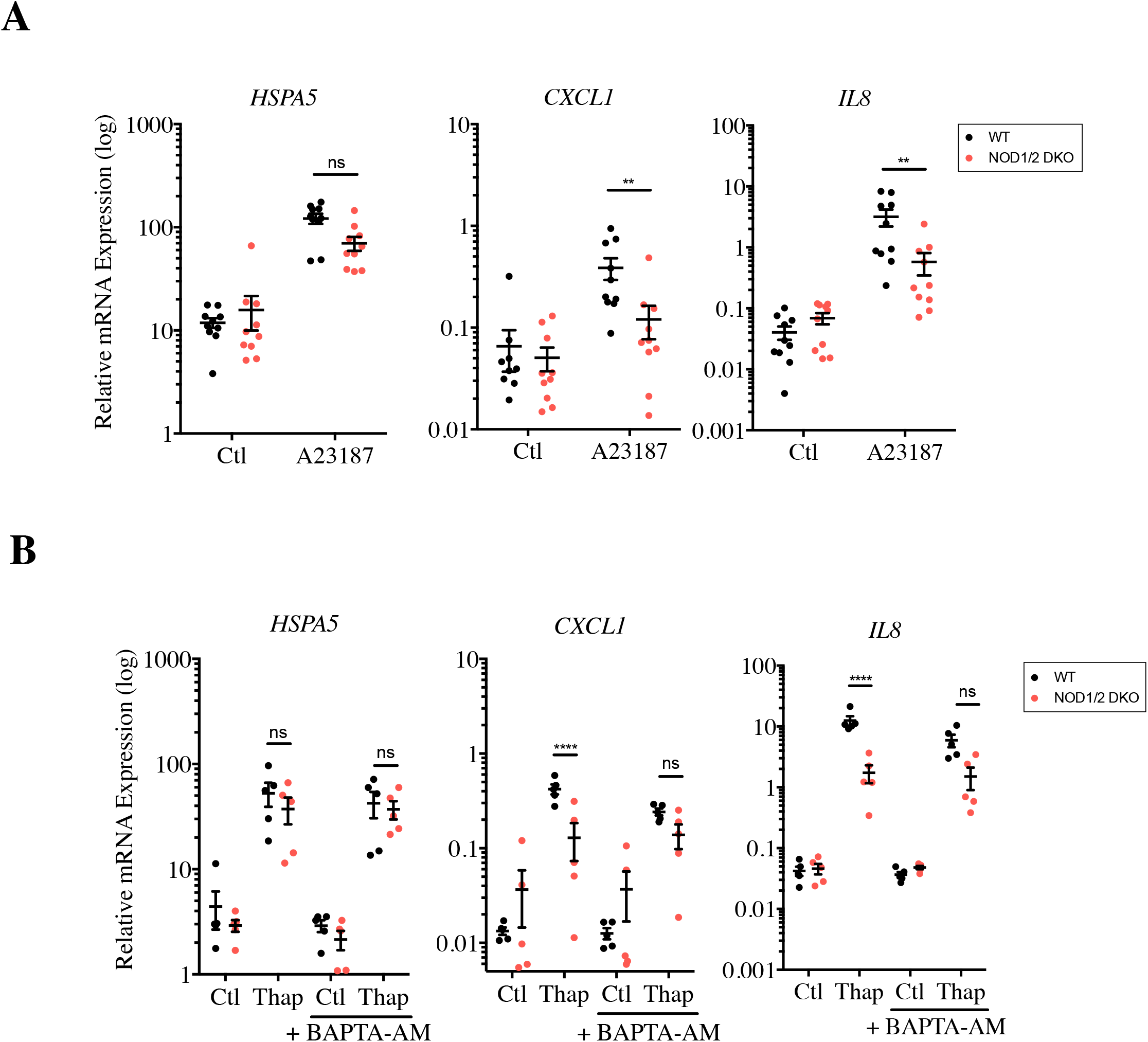
An increase in intracellular Ca^2+^ levels induces NOD-dependent pro-inflammatory signalling. **(A-B)** Expression of *HSPA5, CXCL1* and *IL8* in wild type (WT) and NOD1/2 double KO (DKO) HCT116 cells following stimulation with 1*μ*g/ml A23187 **(A)** or 0.1*μ*g/ml thapsigargin in the presence or absence of 15 μM BAPTA-AM, for 4 hours **(B)** and measured by qPCR. Each point is from an independent experiment and is the average of three technical replicates. ** and ****, p < 0.01 and 0.0001, respectively. NS, not significant.

We next aimed to delineate the mechanism by which an increase in intracellular Ca^2+^ levels causes NOD-dependent pro-inflammatory signalling. We reasoned that since Ca^2+^ regulates vesicular trafficking, exocytosis and recycling of lysosomes (24,25), it could thus in turn impact on the dynamic rate at which cells perform endocytosis. This led us to speculate that the Ca^2+^-dependent signal that triggers NOD-dependent activation in thapsigargin-stimulated cells could be coming from a factor within the extracellular milieu and be brought in by endocytosis. To test this hypothesis, WT and NOD1/2 DKO cells were grown in synthetic Hanks’ Balanced Salt Solution (HBSS) medium supplemented with Ca^2+^, in the presence or absence of 10% fetal calf serum (FCS) and were then stimulated with thapsigargin. Interestingly, NOD-dependent induction of IL8 following thapsigargin stimulation was only observed in the presence of serum, as thapsigargin was unable to trigger IL8 expression when cells were passaged in a medium containing only HBSS + Ca^2+^ prior to thapsigargin stimulation (Fig. 4A). This effect was Ca^2+^-dependent, since it was lost when serum was added to cells grown in HBSS - Ca^2+^ (Fig. S3A). Finally, we boiled the FCS and filtered it to remove molecules with a molecular weight > 3kDa. This boiled/filtered FCS was still able to potentiate NOD-dependent stimulation of IL8 following stimulation (Fig. S3B). This suggests that a heat-resistant small molecule present in FCS drives Ca^2+^-dependent stimulation of NOD1/2 proteins by thapsigargin.

**Figure 4.**
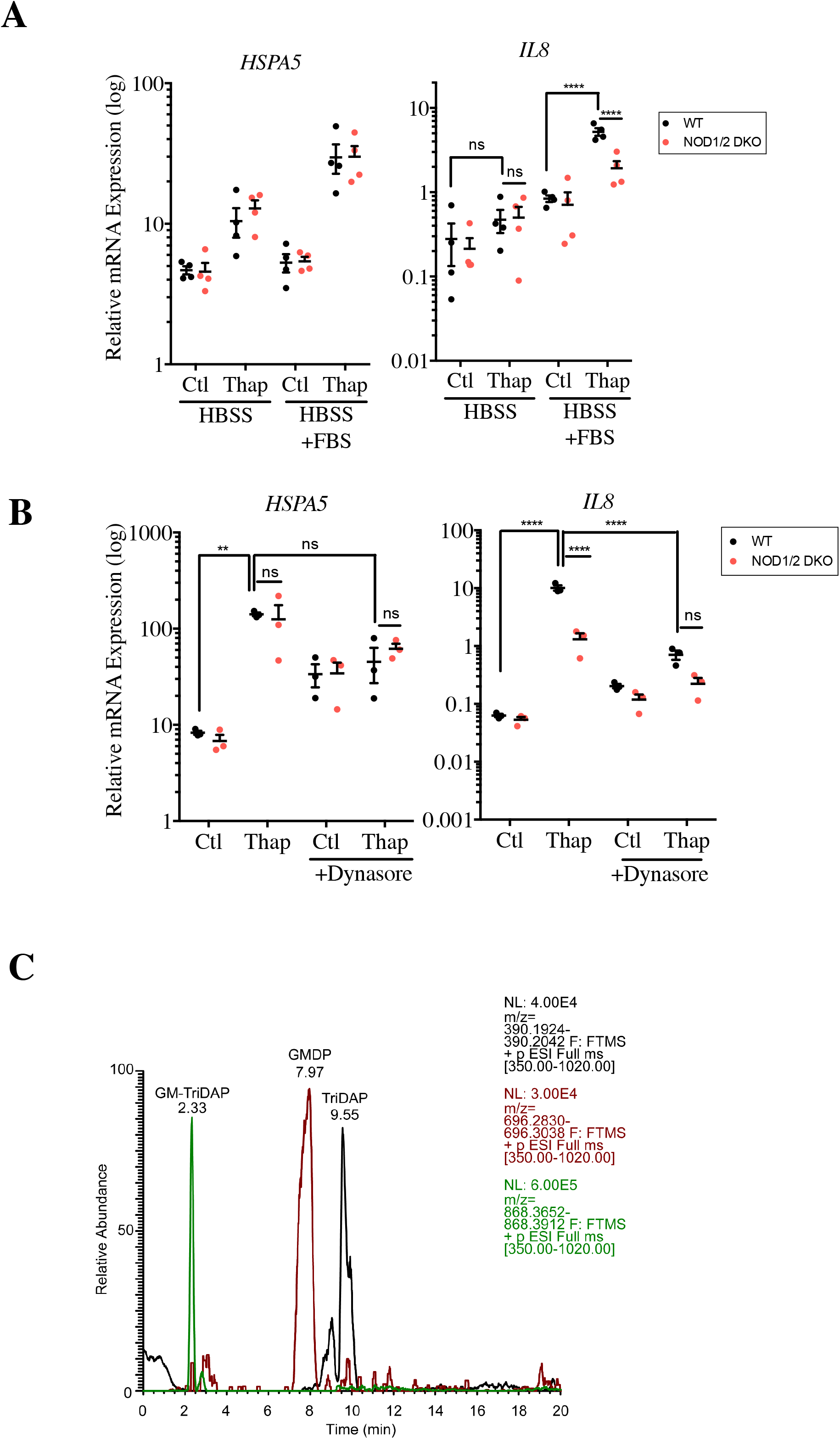
Endocytosis of small molecules from cell culture serum, which contains peptidoglycan fragments, triggers NOD-dependent activation of pro-inflammatory signalling induced by thapsigargin. **(A)** Expression of *HSPA5* and *IL8* in wild type (WT) and NOD1/2 double KO (DKO) HCT116 cells incubated overnight in Hanks’ Balanced Salt Solution (HBSS) medium supplemented with 140 μg/ml CaCl2 in the presence or absence of 10% fecal calf serum (FCS) following stimulation with 0.1*μ*g/ml thapsigargin for 4 hours measured by qPCR. **(B)** Expression of *HSPA5* and *IL8* in wild type (WT) and NOD1/2 double KO (DKO) HCT116 cells following stimulation with 0.1*μ* g/ml thapsigargin, in the presence or absence of 80 mM Dynasore, for 4 hours measured by qPCR. C, LC-MS peak profile of 3 peptidoglycan fragments known to activate NOD1 and NOD2 (GM-TriDAP, GMDP and TriDAP) identified in FCS. Each point is from an independent experiment and is the average of three technical replicates. ** and ****, p < 0.01 and 0.0001, respectively. NS, not significant.

Since internalization of extracellular peptidoglycan into epithelial cells and subsequent activation of NOD1 and NOD2 occurs by clathrin- and dynamin-dependent endocytosis (26,27), we next aimed to define if the active molecule(s) in FCS needed to be internalized by endocytosis to drive thapsigargin-mediated stimulation of NOD proteins. To do so, WT and NOD1/2 DKO HCT116 cells were grown in a normal medium supplemented with 10% FCS and stimulated with thapsigargin in the presence or absence of dynasore, which specifically inhibits endocytosis. We observed that dynasore treatment potently decreased IL8 expression induced by thapsigargin and abolished the NOD dependency of this stimulation (Fig. 4B), implying that the effect required endocytosis. We concluded that a heat-resistant small molecule present in FCS is brought in by endocytosis to stimulate NOD1/2 proteins during thapsigargin stimulation.

All the experiments above point to likely presence of peptidoglycan contaminants in the cell culture grade FCS in which our cells were grown, and that these contaminants would be internalized by endocytosis at a higher rate following an increase in intracellular Ca^2+^, resulting in NOD-dependent stimulation of pro-inflammatory signalling. A previous study similarly reported that peptidoglycan traces found in human serum were responsible for hyphal growth of *Candida albicans* (19). To directly determine if our cell culture FCS contained peptidoglycan fragments, high-pressure liquid chromatography coupled to mass spectrometry was conducted on two separate batches of FCS, which revealed the presence of multiple muramyl peptides and peptidoglycan derived peptides (Fig. S4). In particular, several muramyl peptides identified, including GlcNAc-MurNAc-L-Ala-D-Gln-*meso*DAP (GM-TriDAP), GlcNAc-MurNAc-L-Ala-D-Gln (GMDP) and L-Ala-D-Gln-*meso*DAP (TriDAP) (Fig. 4C), are known activators of NOD1/2 (3). Similar results were obtained when sera from laboratory mice housed in our facility were tested (*data not shown*), suggesting that traces of circulating peptidoglycan in serum may be a common feature.

There are several direct implications of our observations: first, our results provide an alternative explanation for the previous implication of NOD proteins as mediators of inflammatory signalling in response to ER stress (18), by showing that an increase in intracellular Ca^2+^ levels, instead of ER stress itself, caused internalization of trace contaminants of peptidoglycan from the cell culture serum; second, our results suggest that caution should be taken when analyzing if a given stimulus induces NF-κB-dependent pro-inflammatory signalling in a NOD-dependent manner, since multiple pathways can cause transient increase in intracellular Ca^2+^ levels, which could in turn trigger internalization of peptidoglycan contaminants.

More generally, the presence of trace elements of peptidoglycan in animal serum may be of fundamental importance for physiology, although its implication has not yet been carefully evaluated. Interestingly, early studies have demonstrated that circulating peptidoglycan-derived muramyl peptides were able to regulate slow-wave sleep in rabbits (28–30). More recently, it was shown that translocation of peptidoglycan fragments from the intestinal microbiota into the circulation induced functional priming of neutrophils in a NOD1-dependent manner in mice (31), suggesting that these peptidoglycan fragments were sufficient to enhance systemic innate immunity. Since systemic administration of muramyl peptides has been shown to have numerous physiological consequences, from boosting innate and adaptive immune responses (32–34) to triggering protection against obesity-induced insulin tolerance (35) in the case of NOD2 agonists (and opposite effects in the case of NOD1 activators (35,36)), it is tempting to speculate that tonic low-level activation of NOD1/2 systemically caused by circulating peptidoglycan fragments may have far-reaching implications for host physiology that need to be fully characterized.

## Acknowledgements

We wish to thank James and Adrienne Paton (University of Adelaide, Australia) for the kind gift of the SubAB toxins. This work has been supported by grants from the Canadian Institutes for Health Research (CIHR) for the laboratories of DJP and SEG.

## Experimental Procedures

### Cell culture

The human epithelial HCT116 cell line (American Type Culture Collection) was cultured in Dulbecco’s modified Eagle medium (DMEM) supplemented with 10% fetal calf serum (FCS), 2 mM L-glutamine, 50 IU penicillin, and 50 μg/ml streptomycin (Wisent Bio Products). Cells were maintained in 95% air, 5% CO2 at 37°C. Endotoxin-free FCS and phosphate-buffered saline (PBS) were from Wisent (Saint-Bruno-de-Montarville, Quebec, Canada). In some experiments, cells were washed with PBS and cell culture medium was replaced with Hank’s Balanced Salt Solution (Thermo Fisher) supplemented or not with CaCl2 (Sigma; 140 μg/ml).

### Reagents

Muramyl dipeptide (MDP) and iE-DAP (both at 10 μg/ml) were from Invivogen. Thapsigargin (0.1 μg/ml), tunicamycin (1 μg/ml), A23187 (1 μM), dynasore monohydrate (80 μM) and BAPTA-AM (15 μM) were from Sigma. SubAB and SubA272B (50 ng/ml) were a kind gift from Drs James and Adrienne Paton (University of Adelaide, Australia).

### RNA isolation and quantitative RT-PCR

RNA samples were prepared using the GeneJET^™^ RNA Purification Kit (Thermo Scientific) according to the manufacture’s protocol. Eluted RNA was treated with DNase I (Fermentas) at 37°C for 1 h to remove genomic DNA. cDNA was prepared from 1 μg of total RNA using OligoDT, random hexamers, dNTPs, RNase OUT (Invitrogen) and MMLV Reverse Transcriptase (Sigma). cDNA was diluted accordingly and prepared in 12 μL reactions using SYBR^®^ Green qPCR Mastermix (Applied Biosystems). The CFX384 TouchTM Real-Time PCR Detection System (BioRad) was used to obtain the raw CT values. Results were analyzed using the 2^-ΔCt^ formula normalizing target gene expression to the TBP housekeeping control.

### Primary murine organoids

To generate organoid cultures, crypts from the small intestines of mice were extracted as previously described (37). Briefly, the villi of the small intestine were removed by scraping, followed by washing with cold PBS. The remaining tissue were then homogenized and incubated in 2 mM EDTA in PBS for 30 min at 4 °C, followed by vigorous washing in PBS several times to obtain crypt-enriched supernatant fractions. The supernatant fractions were then passed through a cell strainer, pelleted at 300 g for 5 min at 4 °C and resuspended in 50 ul Matrigel (Corning). The crypt-containing organoids were cultured by plating onto the center of a 24-well plate and grown in 500 ul crypt culture medium supplemented with growth factors (R-spondin 1, Noggin, EGF). Organoids were allowed to grow 7 days, followed by passaging onto 6-well plates for stimulation (roughly 10-15 isolated organoids).

### NF-κB luciferase assay

To measure NF-κB luciferase activity, 250,000 HCT116 cells (WT or RIP2 KO) were plated per well and transfected with beta-galactosidase, NF-κB luciferase reporter and pcDNA3 plasmids for 24 h. 10 micrograms of Muramyl dipeptide (MDP) or iE-DAP were added directly to the cell culture medium at the time of transfection, as previously (3). Cells were then gently washed with PBS and lysed in luciferase buffer, followed by incubation at room temperature for 10 min. 10 μL of each cell lysate was then added to a black 96-well plate along with 100 μL of luciferin buffer and luminescence was read using the Victor^3^ plate reader. To measure expression of the beta-galactosidase construct (transfection control) for normalization, 10 μL of original cell lysates were added to 100 μL of the an ONPG buffer, incubated at 37° for 30 min and luminescence was read. The beta-galactosidase values were then used to normalize the absorbance values obtained for each sample.

### LC/MS analysis of serum samples

Aliquots of FCS from different lots were tested as well as serum from laboratory mice kept in our facility in specific pathogen free (SPF) conditions. 20 μl of sera were loaded into the LC-MS platform. The LC-MS platform consisted of an Ultimate 3000 UHPLC coupled to a Q-Exactive mass spectrometer equipped with a HESI II source (Thermo Scientific). Control of the system was performed using Thermo XCalibur 2.2 software and Chromeleon 7.2 software, with data processing conducted using Thermo Scientific Quan Browser. Separation by liquid chromatography was conducted on a Thermo Scientific Hypersil Gold C18 column (50mm×2.1 mm, 1.9 μm particle size). The pump was run at a flow rate of 300 μL/min. Solvent A was water containing 0.1% formic acid; solvent B was acetonitrile containing 0.1% formic acid. The gradient was: 0 min, 5% B; 1 min, 5% B; 2 min, 30% B; 3 min, 30% B; 4 min, 50% B; 7 min, 80% B; 9 min, 80% B; 10 min, 98% B; 11 min 98% B; 12 min, 5% B; 18 min, 5% B. Autosampler temperature was maintained at 10 °C and injection volume was 20μL. Data collection was done in positive ionization mode with MS1 scan range m/z 350–1200, resolution 70,000, AGC target of 3e6 and a maximum injection time of 100 ms, MS2 data was collected using a TOP5 method, 0.4 m/z isolation window, 30 NCE, 17,500 resolution, AGC target 1e5 and a maximum injection time of 50 ms. Data collected were analyzed using Quan Browser.

### Statistical analysis

Significant differences between mean values were evaluated using a two-way ANOVA with multiple comparisons using Prism 5.0. In all RT-qPCR experiments presented in this study, each point represents the average (from three technical replicates) from one experiment. Data from at least three independent experiments were pooled to generate each of the graphs presented. (****p < 0.0001, ***p < 0.001, **p < 0.01, *p < 0.05).

**Table 1.**
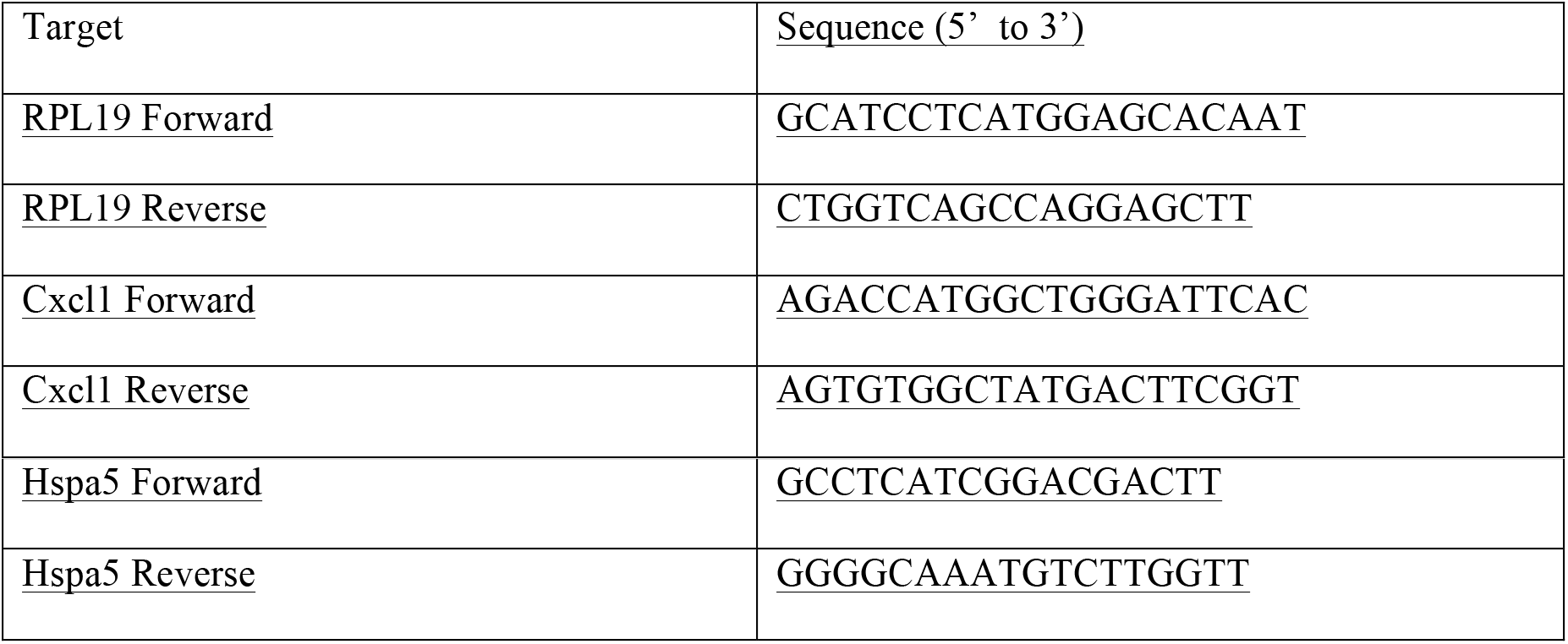
Murine Primers used in this study for qPCR

**Table 2.**
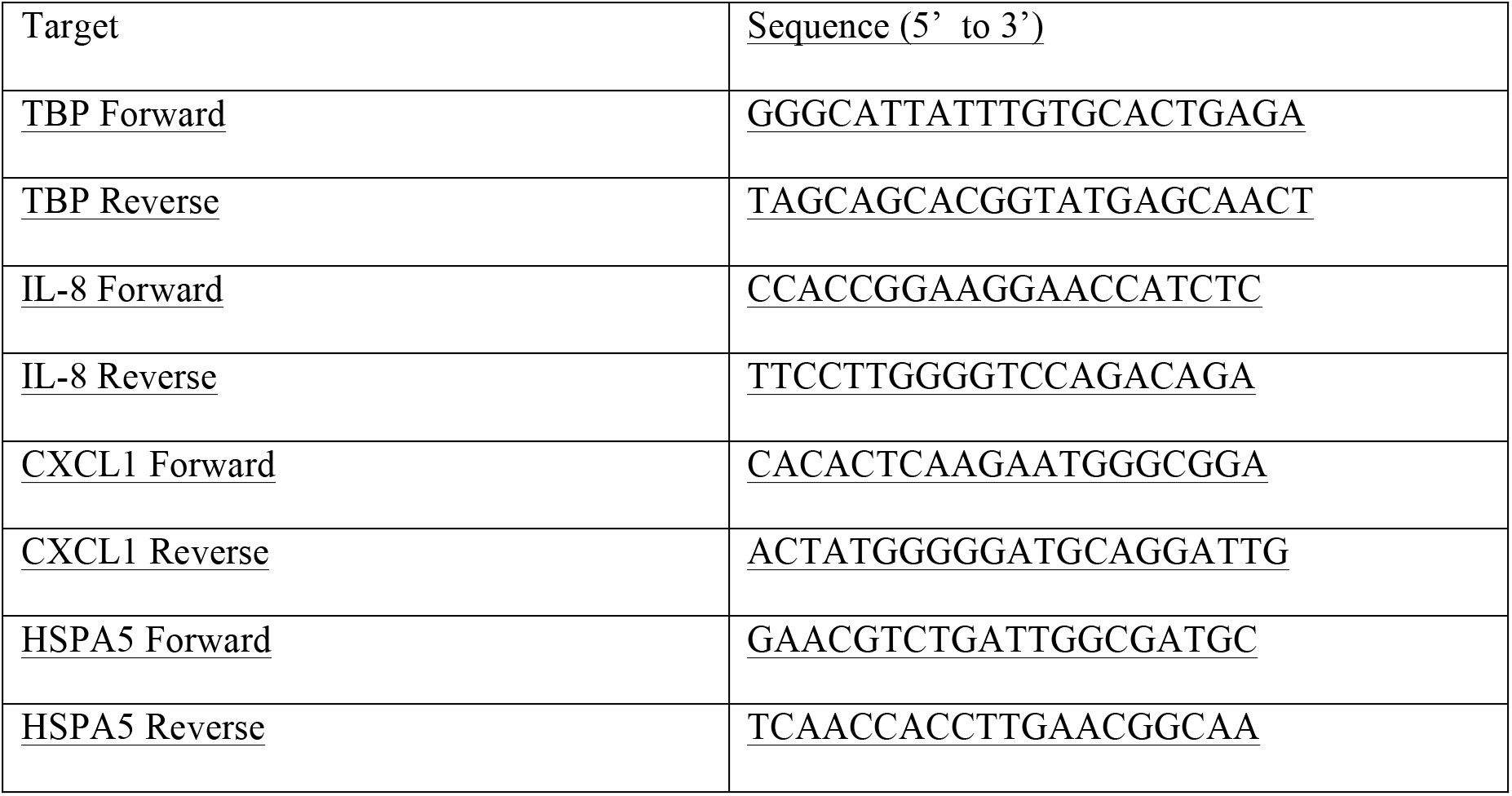
Human Primers used in this study for qPCR

## Supplementary Figure Legends

**Figure S1. Thapsigargin induces pro-inflammatory cytokine expression in a NOD-dependent manner**

**(A-B)** Expression of *IL8* and *CCL20* in wild type (WT), NOD1 knockout (KO), NOD2 KO and NOD1/2 double KO (DKO) HCT116 cells **(A)** or WT and RIP2 KO HCT116 cells **(B)** following stimulation with 0.1*μ*g/ml thapsigargin for 4 hours measured by qPCR. Each point is from an independent experiment and is the average of three technical replicates. *, ** and ***, p < 0.05, 0.01 and 0.001, respectively. NS, not significant.

**Figure S2. Analysis of distinct clones of CRISPR KO HCT116 cells**

**(A)** Wild type (WT) or two distinct clones of RIP2 KO HCT116 cells were transfected overnight with Igκ-Luci (NF-κB reporter construct) and β-galactosidase expressing construct in the presence or absence of 10 μg/ml NOD ligands muramyl dipeptide (MDP) or iE-DAP. **(B)** Expression of *HSPA5* and *CXCL1* in wild type (WT) and two clones of NOD1/2 double KO (DKO) HCT116 cells following stimulation with 0.1*μ*g/ml thapsigargin for 4 hours measured by qPCR. **(C)** Expression of *HSPA5, IL8* and *CCL20* in wild type (WT) and two clones of RIP2 KO HCT116 cells following stimulation with 0.1*μ*g/ml thapsigargin for 4 hours measured by qPCR. Each point is from an independent experiment and is the average of three technical replicates. *, *** and ****, p < 0.05, 0.001 and 0.0001, respectively. NS, not significant.

**Figure S3. Heat-resistant small molecules from cell culture serum trigger NOD-dependent activation of pro-inflammatory signalling induced by thapsigargin**

**(A)** Expression of *HSPA5* and *IL8* in wild type (WT) and NOD1/2 double KO (DKO) HCT116 cells incubated overnight in Hanks’ Balanced Salt Solution (HBSS) medium without Ca^2+^ in the presence or absence of 10% fecal calf serum (FCS) following stimulation with 0.1*μ*g/ml thapsigargin for 4 hours measured by qPCR. **(B)** Expression of *HSPA5* and *IL8* in wild type (WT) and NOD1/2 double KO (DKO) HCT116 cells incubated in DMEM supplemented with either normal 10% FCS (whole) or serum boiled for 10 minutes and filtered with a 3 kDa filter, following stimulation with 0.1*μ*g/ml thapsigargin for 4 hours measured by qPCR. Each point is from an independent experiment and is the average of three technical replicates. *, **, *** and ****, p < 0.05, 0.01, 0.001 and 0.0001, respectively. NS, not significant.

**Figure S4. Analysis of peptidoglycan fragments in two fetal calf serum samples by LC-MS**

List of the peptidoglycan fragments analyzed by LC-MS from two independent fetal calf serum samples. RT, retention time.

