## Supplementary Figures for "Trace levels of peptidoglycan in serum underlie the NOD-dependent cytokine response to endoplasmic reticulum stress"

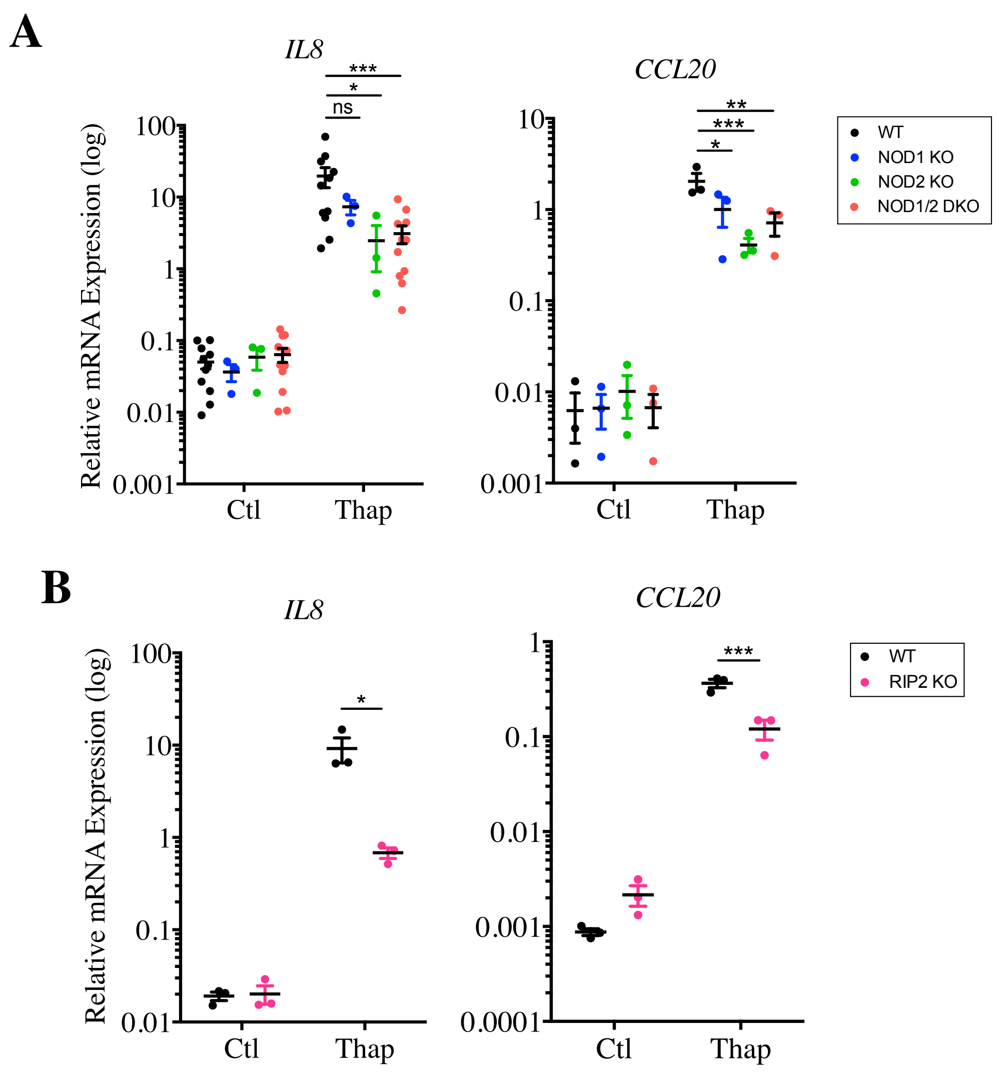
A
IL8
CCL20
Relative mRNA Expression (log)
100
10
1
0.1
0.01
0.001
Ctl
Thap
ns
*
***
**
WT
NOD1 KO
NOD2 KO
NOD1/2 DKO
B
0.0001
RIP2 KO

Figure S1

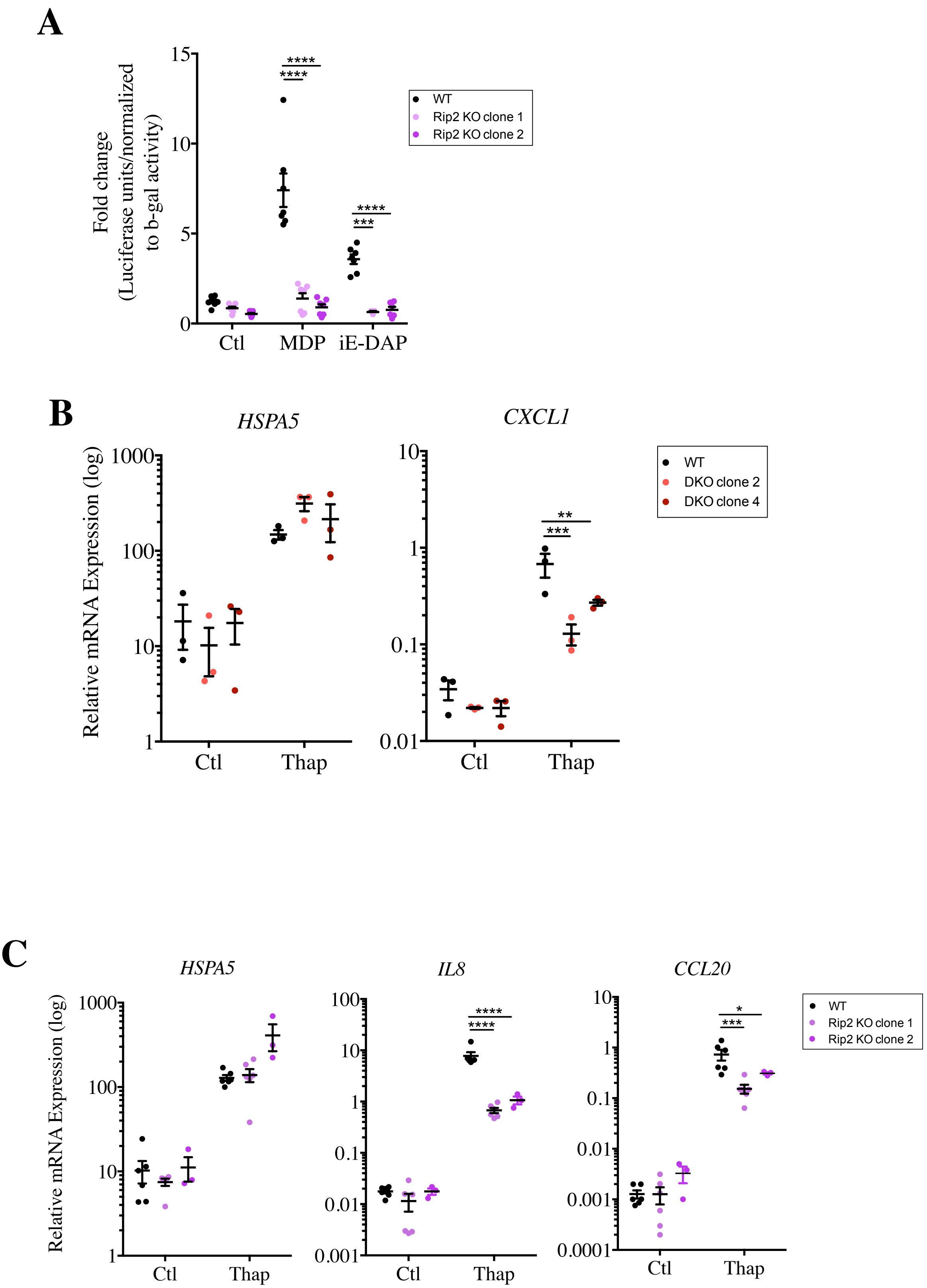
A
Fold change
(Luciferase units/normalized
to b-gal activity)
15
10
5
0
****
****
****
***
Ctl
MDP
iE-DAP
WT
Rip2 KO clone 1
Rip2 KO clone 2
B
HSPA5
CXCL1
Relative mRNA Expression (log)
1000
100
10
1
10
1
0.1
0.01
**
***
Ctl
Thap
WT
DKO clone 2
DKO clone 4
C
HSPA5
IL8
CCL20
Relative mRNA Expression (log)
1000
100
10
1
100
10
1
0.1
0.01
0.001
10
1
0.1
0.01
0.001
0.0001
****
****
*
***
Ctl
Thap
WT
Rip2 KO clone 1
Rip2 KO clone 2

Figure S2

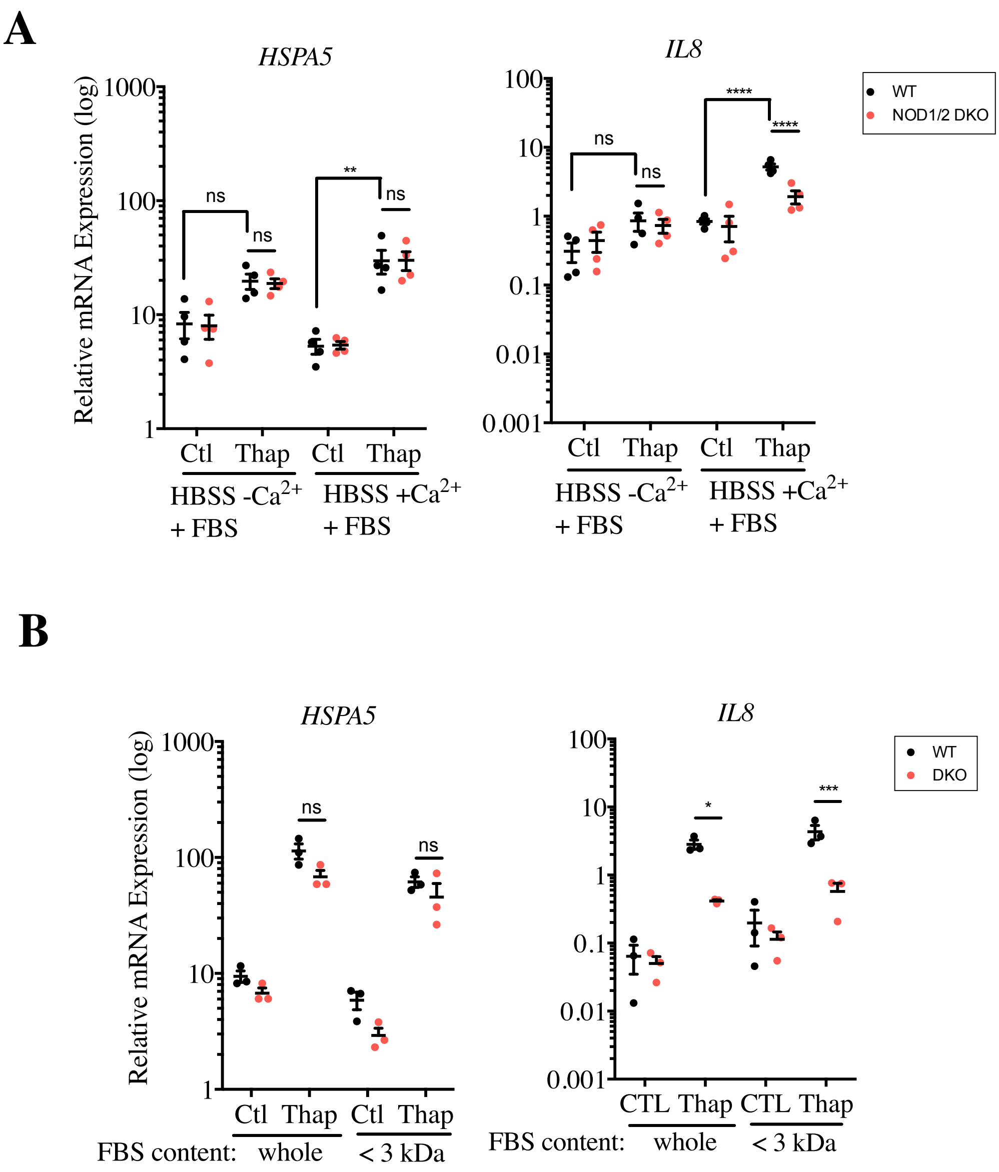
A
HSPA5
Relative mRNA Expression (log)
1000
100
10
1
ns
ns
**
ns
Ctl
Thap
Ctl
Thap
HBSS -Ca2+
+ FBS
HBSS +Ca2+
+ FBS
IL8
100
10
1
0.1
0.01
0.001
ns
ns
****
****
WT
NOD1/2 DKO
B
HSPA5
Relative mRNA Expression (log)
1000
100
10
1
ns
ns
Ctl
Thap
Ctl
Thap
FBS content:
whole
< 3 kDa
IL8
100
10
1
0.1
0.01
0.001
*
***
CTL
Thap
CTL
Thap
FBS content:
whole
< 3 kDa
WT
DKO

Figure S3

| COMPOUND | NEUTRAL FORMULA | M+H | RT | SAMPLE #A | SAMPLE #B |
|---|---|---|---|---|---|
| L-ALA-Γ-D-GLN-MDAP | C15H27N5O7 | 390.1983 | 9.57 | 1.00E+06 | 1.76E+05 |
| L-ALA-Γ-D-GLU-MDAP | C15H26N4O8 | 391.1823 | 14.56 | | 4.27E+04 |
| | | | 10.68 | 1.84E+06 | 1.39E+05 |
| L-ALA-D-GLN-LYS-D-ALA | C17H32N6O6 | 417.2456 | 8.53 | 7.51E+06 | |
| L-ALA-D-GLU-LYS-D-ALA | C17H31N5O7 | 418.2296 | | | |
| L-ALA-D-GLU-MESODAP-D-ALA | C18H31N5O9 | 462.2194 | 10.81 | | 2.05E+05 |
| ANHYDROMURNAC-L-ALA-D-GLN | C19H30N4O10 | 475.2034 | 10.16 | 1.10E+05 | |
| ANHYDROMURNAC-L-ALA-D-GLU | C19H29N3O11 | 476.1874 | 10.9 | 4.92E+05 | 2.05E+05 |
| L-ALA-D-GLN-LYS-D-ALA-D-ALA | C20H37N7O7 | 488.2827 | | | |
| L-ALA-D-GLU-LYS-D-ALA-D-ALA | C20H36N6O8 | 489.2667 | 9.26 | | 1.01E+06 |
| MURNAC-L-ALA-D-GLN | C19H32N4O11 | 493.2140 | | | |
| MURNAC-L-ALA-D-GLU | C19H31N3O12 | 494.1980 | | | |
| L-ALA-D-GLN-MESODAP-D-ALA-D-ALA | C21H37N7O9 | 532.2725 | 2.33 | 1.46E+07 | 5.20E+07 |
| L-ALA-D-GLU-MESODAP-D-ALA-D-ALA | C21H36N6O10 | 533.2565 | | | |
| | | | 9.02 | 5.10E+05 | |
| ANHYDROMURNAC-L-ALA-D-GLU-LYS | C25H41N5O12 | 604.2824 | | | 6.95E+04 |
| MURNAC-L-ALA-D-GLU-LYS | C25H43N5O13 | 622.2930 | | | 1.62E+05 |
| MURNAC-L-ALA-D-GLN-MDAP | C26H44N6O14 | 665.2988 | | | |
| GLCNAC-ANHYDROMURNAC-L-ALA-D-GLU | C27H42N4O16 | 679.2669 | | | |
| GLCNAC-MURNAC-L-ALA-D-GLN | C27H45N5O16 | 696.2934 | 8.03 | 1.02E+06 | |
| GLCNAC-MURNAC-L-ALA-D-GLU | C27H44N4O17 | 697.2774 | | | |
| GLCNAC-ANHYDROMURNAC-L-ALA-D-GLN-MDAP | C34H55N7O18 | 850.3676 | | | |
| GLCNAC-MURNAC-L-ALA-D-GLN-MDAP | C34H57N7O19 | 868.3782 | 2.32 | 5.52E+06 | 4.87E+07 |
| GLCNAC-ANHYDROMURNAC-L-ALA-D-GLU-LYS-L-ALA | C36H59N7O18 | 878.3989 | 1.41 | 7.25E+05 | |
| GLCNAC-MURNAC-L-ALA-D-GLN-LYS-L-ALA | C36H62N8O18 | 895.4255 | | | |
| GLCNAC-ANHYDROMURNAC-L-ALA-D-GLN-MDAP-L-ALA | C37H60N8O19 | 921.4047 | | | |
| GLCNAC-ANHYDROMURNAC-L-ALA-D-GLU-MDAP-L-ALA | C37H59N7O20 | 922.3888 | | | 6.63E+05 |
| GLCNAC-MURNAC-L-ALA-D-GLN-LYS-L-ALA-L-ALA | C39H67N9O19 | 966.4626 | | | |
| GLCNAC-MURNAC-L-ALA-D-GLU-LYS-L-ALA-L-ALA | C39H66N8O20 | 967.4466 | | | 2.42E+06 |
| GlcNac-anhydroMurNac-L-Ala-D-Gln-mDAP-L-Ala-L-Ala | C40H65N9O20 | 992.4419 | | | |
| GlcNac-anhydroMurNac-L-Ala-D-Glu-mDAP-L-Ala-L-Ala | C40H64N8O21 | 993.4259 | | | |
| GlcNac-MurNac-L-Ala-D-Gln-mDAP-L-Ala-L-Ala | C40H67N9O21 | 1010.4524 | | | |
| GlcNac-MurNac-L-Ala-D-Glu-mDAP-L-Ala-L-Ala | C40H66N8O22 | 1011.4364 | | | |

Figure S4
